# A Comparison of Quantitative Mass Spectrometric Methods for Drug Target Identification by Thermal Proteome Profiling

**DOI:** 10.1101/2023.02.15.528618

**Authors:** Amy L. George, Frances R. Sidgwick, Jessica E. Watt, Mathew P. Martin, Matthias Trost, José Luis Marín-Rubio, Maria Emilia Dueñas

## Abstract

Thermal proteome profiling (TPP) provides a powerful approach to studying proteome-wide interactions of small therapeutic molecules and their target and off-target proteins, complementing phenotypic-based drug screens. Detecting differences in thermal stability due to target engagement requires high quantitative accuracy and consistent detection. Isobaric tandem mass tags (TMT) are used to multiplex samples and increase quantification precision in TPP analysis by data-dependent acquisition (DDA). However, advances in data-independent acquisition (DIA) can provide higher sensitivity and protein coverage with reduced costs and sample preparation steps. Herein, we explored the performance of different DIA-based label-free quantification (LFQ) approaches compared to TMT-DDA for thermal shift quantitation. Acute myeloid leukaemia (AML) cells were treated with losmapimod, a known inhibitor of MAPK14 (p38α). Label-free DIA approaches, and particularly the library-free mode in DIA-NN, were comparable or better than TMT-DDA in their ability to reproducibly detect target engagement of losmapimod with MAPK14 and one of its downstream targets, MAPKAPK3. Using DIA for thermal shift quantitation is a cost-effective alternative to labelled quantitation in the TPP pipeline.

## Introduction

Target deconvolution is a key step in the drug discovery pipeline for validating compound-target engagement, determining the mechanism of action, and probing interactions with unexpected proteins to identify possible new therapeutic targets and off-target toxicities.^1^ Thermal proteome profiling (TPP) provides a powerful approach for studying proteome-wide interactions of therapeutic molecules and their target or off-target proteins.^2^ It combines the principles of the cellular thermal shift assay (CETSA) with multiplexed quantitative mass spectrometry, as direct drug binding can elicit conformational changes in a protein, affecting its thermal stability.^3,4^ Cell extracts, which lack normal cellular metabolism, are used to investigate thermal shifts due to direct compound engagement, although proteins are removed from their native environment, which may also affect thermal stability.^2, 5, 6^ Treatment of intact cells or tissues allows additional understanding of cellular response in a physiologically relevant setting, impacting protein thermal stability while capturing downstream effects.^2^ The method has been applied to interrogate drug-target interaction, protein–substrate interaction, protein degradation, and post-translational modifications (PTMs) in various biological systems.^7–10^

Quantifying differences in thermal stability requires high quantitative accuracy and consistent detection. Isobaric labelling with tandem mass tags (TMT) has been traditionally used with data-dependant acquisition (DDA) to quantify proteome-wide changes in thermal stability as the pooling of samples increases throughput, reduces technical variability and is suitable for fractionation thus providing deep proteomic coverage.^11^ Initially, ten temperatures labelled with TMT-10plex^12^ were recommended to generate melting curve data points, requiring two TMT experiments per biological replicate, for treatment and vehicle control conditions. ^4^ Since the development of TMTpro 16-plex and 18-plex, performing eight (or nine) temperatures for melting curve generation allows two conditions to be analysed within a single experiment per biological replicate.^13–15^ Sample-specific reporter-ions produced after fragmentation are used for quantification, reducing the number of missing peptide quantification values as peptides across conditions are not separately subject to semi-stochastic precursor sampling as with label-free DDA quantification but are acquired within the same MS^2^ scan.^16^ While this quantitative approach works very well within the boundaries of its multiplicity, principal drawbacks include the high price of reagents, extended sample preparation time and constraints on sample number analyses compared with label-free approaches. In addition, notable increases in missing protein and peptide values are observed once multiple batches are integrated.^17^ Increased variability is particularly seen with the cell-based TPP protocol due to sample-specific lysis, requiring additional replicates for reproducible thermal stability shifts to become significant. ^4^ High workflow costs frequently restrict data collection and therefore many TPP studies were performed with just two biological replicates, which can limit result significance. ^5, 18, 19^ Moreover, quantification of MS^2^ fragments can suffer from ion interference and ratio compression, leading to a dampening of fold changes and under-measure of thermal shifts.^20^,^21^ Synchronous precursor selection (SPS) MS^3^ methods (partly) overcome this by isolating fragments after MS^2^ for further fragmentation, resulting in improved quantitative accuracy, but reduced proteome coverage due to lower scan rates and is limited to Tribrid instruments. ^22^

Label-free quantification (LFQ) acquired in DDA has been used as a less expensive alternative to isobaric methods for thermal shift quantitation for target deconvolution. In the work from Türkowsky *et al.*, the gain in flexibility with LFQ-DDA enabled adaptation of the TPP protocol to investigate oxygen-sensitive proteins in anaerobic bacteria. However, low proteomic sequence coverage and a high proportion of missing values in melting curves were reported, likely due to the intensity-triggered precursor selection biases accompanying DDA quantification.^23^, ^24^ Alternatively, data-independent acquisition (DIA) is a single-shot method that provides precise, accurate proteome quantification of label-free samples with low missing values and high analytical depth, as all detectable peptides within specific *m/z* windows are fragmented and quantified in parallel. ^11,18^ This quantitative approach has previously been adopted for target deconvolution in limited proteolysis workflows to measure stability of protein-ligand complex under proteolytic conditions.^25^ More recently, it was adopted in a matrix thermal shift assay to detect ligand concentration-dependent stabilisation of proteins, at a single melting temperature.^26^ Ruan *et al.* treated K562 lysates with staurosporine, a model widely used for thermal shift assay method evaluation and demonstrated reduction in sample preparation time, cost and effort. By adopting a library-free, DirectDIA approach, they increased throughput and achieved comparable sensitivity for identifying kinase targets to a recent 2D-TPP TMT study. ^13^, ^26^

There are now various pipelines for DIA peptide identification, each with distinct strengths and suitability depending on the type of proteomic data.^27–29^ Project-specific libraries generated by analysing pre-fractionated DDA samples, which are representative of the model system, historically provide the greatest proteome coverage but largely depend on the size and quality of the library, increasing instrument run-time.^27,30,31^ Hybrid libraries can be constructed in Spectronaut to increase library depth and precision by combining the DDA library with DIA data.^32^ However, library-free approaches require no additional data to generate spectral libraries and have been demonstrated to achieve nearly equivalent whole-proteome coverage.^33, 34^ Library-free analysis is commonly performed in Spectronaut using DirectDIA, where Pulsar performs a classical database search directly on the DIA runs to create a library which is then used for a targeted analysis of the same DIA runs.^35^ An alternative approach is to produce an *in silico* spectral library using software such as DIA-NN, which has not yet been trialled for analysing TPP data.^36^

In this study, we explored the performance of different library-free and library-based DIA approaches in a TPP workflow, and benchmarked these with traditional TMT-DDA for thermal shift quantitation. We use losmapimod, a potent MAPK14 ATP competitive inhibitor, to provide a known true positive hit in acute myeloid leukaemia (AML) cells. While all five workflows reliably measured thermal stabilisation of MAPK14 and its downstream effector, MAPKAPK3, library-free mode DIA-NN showed superior performance as a cost-effective alternative to TMT-DDA for thermal proteome profiling.

## Experimental Section

### Compounds

Losmapimod (GW856553) was purchased from Biorbyt (UK); human recombinant IL-1β and TNF-α proteins from PeproTech (UK). A clinicaltrials.gov search on February 13, 2023, for losmapimod (keywords: losmapimod, GW856553, GW856553X, SB856553 or GSK-AHAB).

### Cell culture

THP-1 (ATCC, TIB-202) cell line was cultured in RPMI 1640 (Gibco) supplemented with 10 % heat-inactivated foetal bovine serum and 2 mM L-glutamine at 37°C in a humidified 5 % CO_2_ atmosphere. ATCC routinely performs cell line authentication, using short tandem repeat profiling as a procedure. Cell experimentation was always performed within a period not exceeding six months after resuscitation in mycoplasma-free culture conditions.

### Cell proliferation and viability assay

Cells were pre-treated with 1 μM losmapimod for 1 h before stimulation with 10 ng/mL IL-1β and/or 20 ng/mL TNF-α for 15 min. Cell proliferation was assessed at different time points (3, 5, 7 and 9 days) using AlamarBlue Cell Viability Reagent (Invitrogen) per manufacturer’s instructions. Following 4 h incubation, the plate was read with the SpectraMax iD3 Fluorescence Microplate Reader (Molecular Devices) using 555/595 nm (excitation/emission) filter settings. Trypan Blue solution was used for the determination of cell viability.

### Surface plasmon resonance (SPR)

SPR ligand interactions assays were performed on a Biacore S200 (Cytiva Life Sciences) at 2°C using multi-cycle settings. Biotinylated avidin-MAPK14 protein (MRC-Reagents, Dundee) was immobilized onto a Streptavidin surface chip, through injection of 50 μg/mL MAPK14 in DMSO-free SPR running buffer (20 mM HEPES, 150 mM NaCl, 0.1 mM EGTA, 0.5 mM TCEP, 0.01 % Tween-20, pH 7.4) over the active flow cell eliciting final captured response units (RUs) of 7719 RUs. The inhibitor analytes (20 mM HEPES, 150 mM NaCl, 0.1 mM EGTA, 0.5 mM TCEP, 0.01 % Tween-20, pH 7.4, 1 % DMSO) were then injected over both control and active surfaces for 90 seconds at 30 μL/min before being allowed to dissociate for 600 seconds over ten concentration series to record dose-responses: 0.05 nM – 333.33 nM. A solvent correction was applied to the data collection and used an 8-point DMSO solvent correction was applied. Responses were analyzed using Biacore Evaluation Software (Cytiva Life Sciences) using affinity fit to determine the Kd. Data are representative of three technical replicates.

### Western blot

Ten micrograms of protein from each sample were mixed with 5X Laemmli buffer with 5 % β-mercaptoethanol, heated for 5 min at 100°C, and separated on 10 % SDS-PAGE gels. Following electrophoresis, the proteins were transferred to Immobilon-P transfer membranes (Sigma-Aldrich) using the Trans-Blot^®^ TurboTM Transfer System (Bio-Rad Laboratories). Membranes were visualised with Ponceau Red (FlukaTM Analytical). Membranes were probed with the following antibodies purchased from Cell Signaling Technology: MAPKAPK3 (#7421), MAPK14 (#9218), Thr180/Tyr182-P-p38 MAPK (#4511), anti-rabbit IgG-HRP (#7074). GAPDH (sc-47724) was purchased from Santa Cruz. Amersham Imager 600 digital imaging system (GE Healthcare) was used for image acquisition and Quantity One^®^ v4.6.3 (Bio-Rad Laboratories) for band densitometry.

### Cellular thermal shift assay

Intact-cell TPP experiments were performed on THP-1 cells as described previously, with some slight alterations.^15^ Briefly, thirty million THP-1 cells were treated with 1 μM losmapimod or vehicle (0.01 % DMSO) in complete media at 37°C for 1 h. Cells were collected and washed twice with PBS containing 1 μM losmapimod or vehicle. Washed cells were suspended in PBS supplemented with treatment or vehicle plus 0.4 % NP-40, cOmplete Protease Inhibitor Cocktail (Sigma-Aldrich) and phosphatase inhibitor cocktail (1.2 mM sodium molybdate, 1 mM sodium orthovanadate, 4 mM sodium tartrate dihydrate, and 5 mM glycerophosphate), then separated into 8 fractions for thermal profiling. Fractions were heated at 40, 44, 48, 52, 56, 60, 64, and 68°C for 3 min, incubated for 3 min at room temperature, and snap-frozen at −80°C. Samples were lysed with four freeze-thaw cycles using dry ice and a thermo block at 35°C. Cell lysates were centrifuged at 100,000 x g for 20 min at 4°C to separate protein aggregates from soluble proteins. Supernatants were collected and protein concentrations were determined using Pierce BCA Protein Assay Kit (Thermo Fisher Scientific) for western blot and mass spectrometry analysis.

### Sample preparation for mass spectrometry

#### Reduction, alkylation, and digestion of soluble fractions for quantitative proteomic analysis

Volumes equivalent to 20 μg of protein per sample were made equal to a final concentration of 5 % sodium dodecyl sulfate (SDS) in 50 mM Triethylamonium bicarbonate (TEAB) and 5 mM tris(2-carboxyethyl)phosphine (TCEP, Pierce), incubated for 30 min at 47°C, then alkylated with 10 mM iodoacetamide for 30 min at room temperature in the dark. Protein digestion was performed using the suspension trapping (S-Trap^™^) sample preparation method according to the manufacturer’s guidelines (ProtiFi, USA). Proteins were digested with trypsin (Worthington-Biochem) in 50 mM TEAB pH 8.0 at an enzyme-to-protein ratio of 1:10 (w/w) for 2 h at 47°C. Peptides were eluted by three successive washes of 40 μL 50 mM TEAB pH 8.0, 40 μL 0.2 % formic acid (FA) in water and 35 μL 0.2% FA in 50 % acetonitrile, respectively. The resulting eluates were vortexed and divided into two equal aliquots for LFQ-DIA analysis or TMT preparation, and dried before storage at −80°C.

#### TMTpro-16plex peptide labelling and offline HPLC fractionation

Isobaric labelling of peptides was performed using TMTpro^™^ 16plex Label Reagent Set (Fisher Scientific, UK) according to the manufacturer’s recommended protocol (lot number: WC320807). After confirming labelling efficiency of > 97 % by short 1 h LC-MS runs, labelled peptides from each temperature point were combined to a single sample per biological replicate and desalted with C18 Macro Spin Columns (Harvard Apparatus, USA). The pooled sample was subject to fractionation using basic-pH reversed-phase liquid chromatography (LC) on a Gemini C18 column (250 mm × 3 mm, 3 μm, 110 Å; Phenomenex) on a Dionex Ultimate 3000 off-line LC system, generating 12 fractions which were dried. All solvents used were high-performance liquid chromatography (HPLC) grade (Fisher Scientific). Mobile phase A consisted of 20 mM ammonium formate pH 8.0 and mobile phase B of 100 % acetonitrile (MeCN). Peptides were separated over a 49 min linear gradient of 1 – 49 % B at 250 nL/min. Peptide elution was monitored by UV detection at 214 nm. Each of the twelve fractions were acidified to a final concentration of 1 % trifluoroacetic acid (TFA) and dried using a speed-vac.

#### Sample preparation of THP-1 soluble fraction pool for library generation

To generate DDA data for project-specific libraries, 60 μg of pooled soluble-fraction digest was subject to fractionation as described above for the TMTpro labelled peptides, generating a total of 12 fractions.

### Liquid chromatography

All samples (label-free for DIA analysis, or fractioned pools for TMT experiment or DDA library generation) were solubilised in 2 % MeCN with 0.1 % TFA to 0.2 μg/μL concentration before being injected in volumes equivalent to 1 μg on an UltiMate 3000 RSLC nano System (Thermo Fisher Scientific). Peptides were trapped for 5 min in A (0.1 % FA in water) at a flow rate of 10 μL/min on a PepMap 100 C18 LC trap column (300 μm ID x 5 mm, 5 μm, 100 Å) then separated using an EASY-Spray analytical column (50 cm x 75 μm ID, PepMap C18, 2 μm, 100 Å) (Thermo Fisher Scientific) flowing at 250 nL/min. The column oven temperature was set at 45°C. All peptides were separated using an identical linear gradient of 3 – 35 % B (80 % MeCN containing 0.1 % FA) over 120 min.

### Mass spectrometry

All data was acquired on an Orbitrap Fusion Lumos Tribrid mass spectrometer (Thermo Fisher) in positive ion mode.

#### TMT-labelled samples

Data were acquired in DDA mode with positive ion mode. Full MS spectra (*m/z* 375 – 1,500) were acquired at 120,000 resolution, automated gain control (AGC) target 4 × 10^5^ and a maximum injection time of 50 ms. The most intense precursor ions were isolated with a quadrupole mass filter width of 0.7 and higher-energy collision-induced dissociation (HCD) fragmentation was performed in one-step collision energy of 30 % and 0.25 activation Q.

Detection of MS^2^ fragments was acquired in the linear ion trap in rapid scan mode with a AGC target 1 × 10^4^ and a maximum injection time of 50 ms. An electrospray voltage of 2.0 kV and capillary temperature of 275°C, with no sheath and auxiliary gas flow, was used. Dynamic exclusion of previously acquired precursor was enabled for 60 s with a tolerance of ± 10 ppm. Quantitation of TMT-tagged peptides was performed using FTMS3 acquisition in the Orbitrap mass analyser operated at resolution of 60,000, with a standard AGC target and maximum injection time of 118 ms. HCD fragmentation of MS^2^ fragments in one-step collision energy of 55 % to ensure maximal TMT reporter ion yield and enable synchronous-precursor-selection (SPS) to include ten MS^2^ fragment ions in the FTMS^3^ scan.

#### Label-free samples

Full scan spectra (*m/z* 390 – 1,010) were acquired in centroid mode at an Orbitrap resolution of 60,000, an AGC target set to standard, a maximum injection time of 55 ms, RF lens at 30 % and expected peak width of 20 s. Subsequently, an 8 *m/z* staggered window scheme^24^ was used to collect DIA scans, utilising 75 windows, with a 4 Da window overlap. HCD collision was set to 33 %, loop count of 75, Orbitrap resolution of 15,000, AGC target of 100 % and a maximum injection time of 23 ms.

#### DDA library generation

Data were acquired in DDA with positive ion mode. Full MS spectra (*m/z* 350 – 1,000) were acquired at 120,000 Orbitrap resolution, using standard AGC and a maximum injection time of 50 ms. The most intense precursor ions were isolated with a quadrupole mass filter width of 1.6 *m/z* and HCD fragmentation was performed in one-step collision energy of 30 % and 0.25 activation Q. Detection of MS^2^ fragments was acquired in the linear ion trap in rapid scan mode with a standard AGC target and a maximum injection time set to auto. Dynamic exclusion of previously acquired precursor was enabled for 38 s with a tolerance of ±. 10 ppm.

### Raw data processing

#### TMT-DDA peptide identification

Raw TMT-DDA files were loaded into MaxQuant (v1.6.10.43) and searched against the *Homo sapiens* Uniprot database containing 42,426 entries with isoforms (downloaded February 2021). Specific search parameters included: trypsin as the protease for digestion, a maximum of two missed tryptic cleavage sites per peptide; dynamic modifications included oxidation of methionine and N-terminal acetylation, and fixed modifications included carbamidomethylation of cysteine residues. Reporter ion MS^3^ was used for quantification. Protein identifications were filtered to a false discovery rate (FDR) of less than 1 %, and features matching a contaminant or reverse peptide; only identified by site; or which contained less than two unique peptides were removed. Reporter ion intensity corrected columns were used for downstream data analysis.

#### LFQ-DIA library generation

Spectronaut Pulsar (v16.1, Biognosys, Switzerland) was used to construct spectral libraries. DDA raw files were searched with Pulsar to generate a search archive (DDA library). DIA files were subsequently searched in combination with the DDA search archive to produce a hybrid library.

#### LFQ-DIA peptide Identification

Raw LFQ-DIA files were processed with Spectronaut (v16.1, Biognosys, Switzerland) and analysed either library-free using default setting with DirectDIA, or a library-based search performed using the previously constructed DDA library or hybrid library. Search parameters were identical to those previously specified in MaxQuant search, except MS^2^ was used for quantification. Data filtering was set to Q-value and normalisation set to automatic. All datasets were filtered in Spectronaut to remove single hits, decoys, proteins with less than two unique peptides and an FDR of less than 1 %. The same DIA files were analysed library-free using DIA-NN 1.8.1^36^ using default parameters and the same FASTA as all other searches. Resulting identifications were filtered using R (v4.0.4) to include only proteins with more than two unique peptides and an FDR of less than 1 %.

### Thermal proteome profiling data analysis

Soluble protein intensity values of each dataset were log2 transformed, normalised to their median abundance, and expressed as a ratio to the lowest temperature sample (40°C). Each ratio was then normalised to the mean abundance for the identified top 50 temperature-stable proteins in sample 60°C, as described by Miettinen, T. P. *et al.* ^37^ The temperature-range TPP package was used to perform analysis (https://www.bioconductor.org/packages/release/bioc/html/TPP.html).^38^ Data was filtered to include only those protein groups suitable for curve fitting (> 2 valid fold changes per protein). This fitting was used to determine the melting point (T_m_), which is defined as the temperature at which half of the number of proteins was denatured. The melting point differences (ΔT_m_) were calculated by subtracting the T_m_ values of treated and untreated samples. The sigmoidal melting curves were filtered according to the following criteria: melting curves must reach a relative abundance plateau of < 0.3, and the coefficient of determination (R^2^) must be > 0.8. The significance threshold was set to adjusted p-value < 0.05, with a melting point difference of > 2 °C, and a standard deviation (SD) < 2.

### Data availability

The mass spectrometry proteomics data have been deposited to the ProteomeXchange Consortium via the PRIDE partner repository^28^ with the data set identifier: PXDXXXXX.

### Statistical analysis

Statistical analyses of data were performed in R (v4.0.4) or GraphPad prism (v9.0.2).

## Results and Discussion

### Experimental overview

The activation of the p38 MAPK pathway plays a critical role in orchestrating a range of cellular stresses, growth, and survival of tumour cells. ^39, 40^ p38 MAPK is often increased and/or overactivated in AML blasts and represents an important pharmacological target. ^41–43^ However, selective p38 MAPK inhibitors have limited efficacy and for a variety of clinical indications none have progressed to Phase III.^44^ Losmapimod (known as GW856553 or GSK-AHAB) is a selective, potent, and orally MAPK14 (p38α MAPK) ATP competitive inhibitor^45^ (**Figure 1A**) used in several clinical trials (**Table S1**). Here, we validated that losmapimod binds directly to MAPK14 by surface plasmon resonance (SPR) analysis (**Figure 1B**). In addition, losmapimod had slow dissociation rates (K_off_) which have been associated with better selectivity, lower toxicity, and a broader therapeutic window. ^46^ Losmapimod was also able to reduce MAPK p38 activation in the human monocytic leukaemia cell line, THP-1 (**Figure 1C**) and reduced the proliferation of AML cells without affecting viability (**Figure S1).**

**Figure 1.**
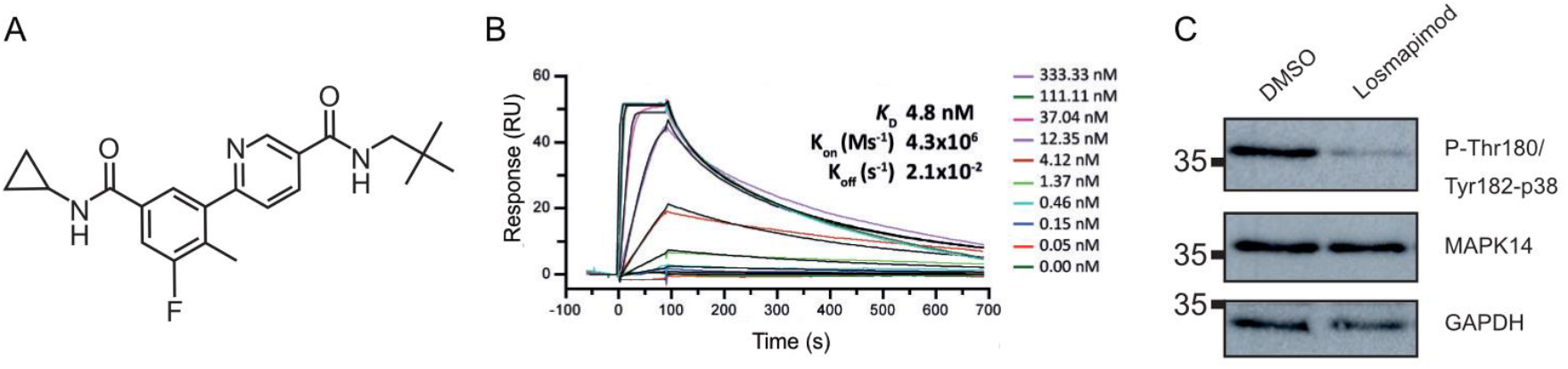
Losmapimod interacts with MAPK14 and reduces p38 MAPK phosphorylation. **A)** Structure of losmapimod.**B)** Assessment of losmapimod binding to MAPK14 using surface plasmon resonance (SPR). Representative multicycle SPR sensograms for losmapimod showing a dose-dependent concentration series against immobilized MAPK14. K_D_, equilibrium dissociation rate constant; K_on_ (Ms^-1^), on-rate constant or association reaction; K_off_ (s^-1^), off-rate constant or dissociation reaction. Experiments were performed in triplicate. **C)** Western blot analysis of MAPK14 and p38 MAPK phosphorylation in THP-1 cells with the vehicle control, DMSO, and 1 μM losmapimod for one hour shows loss of p38 phosphorylation in response to losmapimod. GAPDH serves as a loading control. A representative image of three replicates is shown. Relative mobilities of reference proteins (kDa) are shown on the left of each blot.

To date, losmapimod is the only MAPK14 inhibitor in Phase III clinical trials as it is safe and well-tolerated in previous human clinical studies. Losmapimod therefore represents a valuable therapeutic agent for AML and can provide a positive protein target, MAPK14, for thermal shift analysis and direct method comparison.

As THP-1 cells have basal activation of MAPK14 (**Figure 1C**), cells were treated with the compound or vehicle without stimulus in three independent experiments and incubated at eight temperatures between 40-68°C. Denatured, insoluble protein aggregates were then removed by ultracentrifugation and the protein quantity of each soluble fraction measured (**Figure 2, 1**). Equal protein amounts from the soluble-protein fraction were digested and divided in two to compare the performance of five workflows to detect MAPK14 thermal stabilisation. TMT-DDA is the traditional approach for melting temperature quantitation in a TPP workflow. ^2^ Each temperature for compound and vehicle treated cells were chemically labelled with an isobaric tandem mass tag, using one set of TMTpro 16-plex per replicate. Small aliquots of each sample were pooled, and following clean-up, analysed by DDA-MS to confirm labelling efficiency was <97 %. For each replicate, the 16 peptide samples were pooled into a single peptide mixture, cleaned-up and separated into non-consecutively concatenated fractions by basic-pH reversed-phase LC, producing a total of 12 fractions to be analysed by MS^3^-DDA quantitation (**Figure 2, 2**).

**Figure 2.**
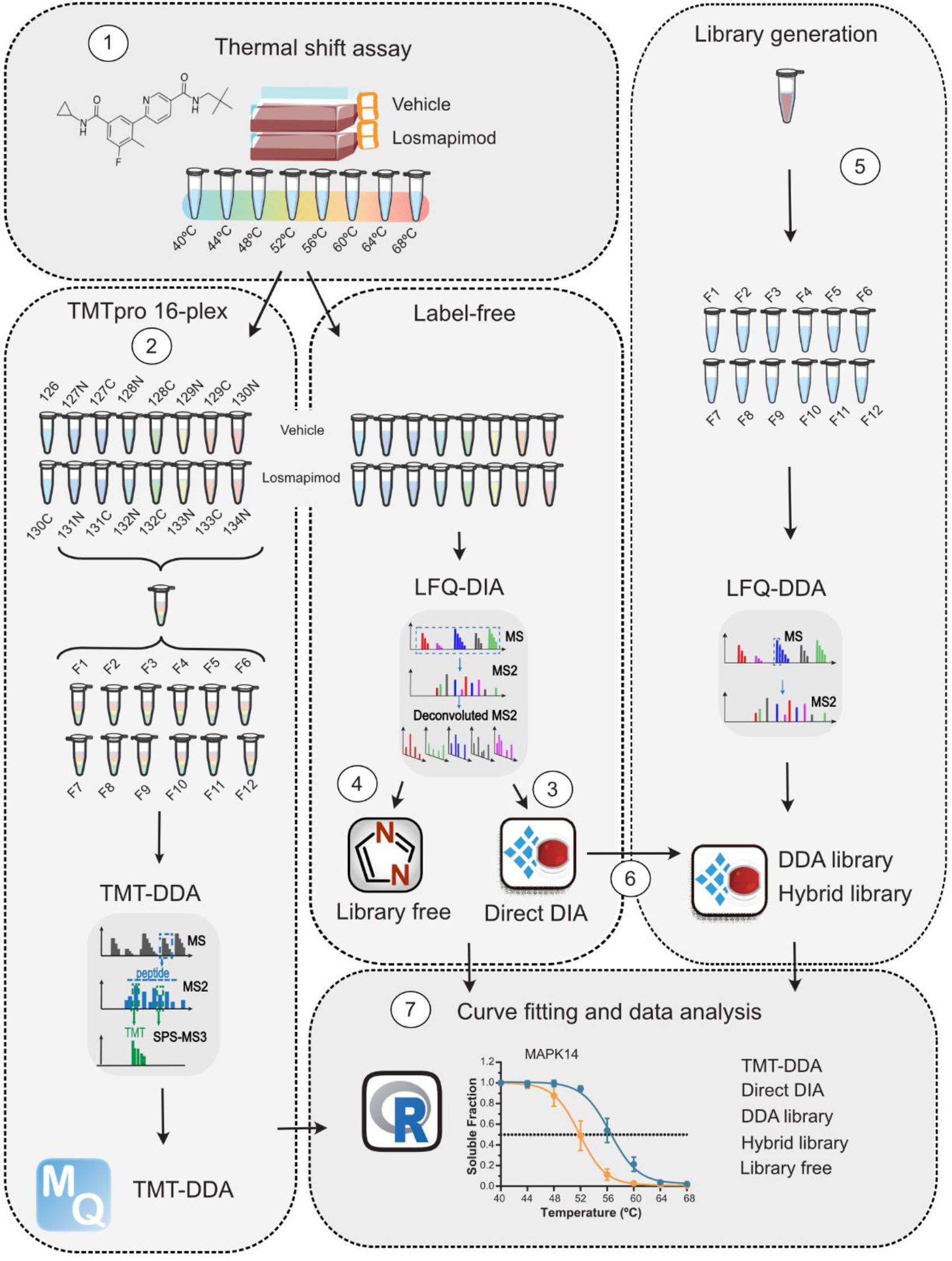
Comparative workflow. **(1)** To provide a benchmark sample with a positive hit for thermal stabilisation, AML cells were treated with 1 μM losmapimod or vehicle for 1 h, then subjected to the thermal shift assay and lysed. Soluble fractions were digested, and each sample split in half for either **(2)** 16-plex TMT-labelling, fractionation, DDA-MS^3^ quantification and analysis in MaxQuant or for label-free DIA analysis using library-free **(3)** DirectDIA with Spectonaut and **(4)** library-free mode DIA-NN or DDA library-based approaches. For library-generation, small volumes from remaining TPP samples were pooled and digested to provide **(5)** a project-specific DDA-based library within Spectronaut, and a **(6)** hybrid library was constructed of both DDA and DIA data. **(7)** After data processing, thermal proteome profiling data analysis was performed in R and the five workflows compared.

The other half of the peptide samples were left unlabelled, analysed by single-shot DIA and processed using four different DIA approaches (DirectDIA, DDA library and hybrid library in Spectronaut, and library-free mode in DIA-NN). Firstly, data was searched using DirectDIA for peptide identification **(Figure 2, 3)**. Based on the acquired DIA data, the software generates a pseudo-MS^2^ library by searching data against a sequence database, which is then employed to analyse the original DIA data. This approach provides advantages of reduced instrument time, efforts and cost by overcoming the need for library generation. ^26^ The data was also searched using library-free mode in DIA-NN which instead generates an *in silico* library from a protein sequence database **(Figure 2, 4)**. ^36^ While their performance have previously been benchmarked and shown unique advantages in large-scale proteomic and phosphoproteomic workflows, they have never been compared for TPP. ^27, 30, 47^

Nevertheless, library-based DIA approaches are still largely implemented to improve depth of proteomic coverage in expression studies^32^ but have also never been explored for thermal shift quantitation. Therefore, the remaining volumes of each soluble fraction were pooled and digested to provide a project-specific reference sample for library generation. Spectral libraries, generated experimentally from peptide fractionation followed by DDA analysis, remain most common in DIA peptide quantification (DDA library).^24,27^ We fractionated the reference sample digest by basic-pH reversed-phase LC, producing 12 fractions to be analysed by MS^2^-DDA quantification. A DDA library was then constructed in Spectronaut which consisted of 69,153 precursors, 58,907 peptides and 7,092 protein groups **(Figure 2, 5).** While DDA library workflows can provide deeper proteome coverage, matrix-effects can influence retention time differences between the fractionated, and consequently less complex, DDA samples compared with quantitative single-shot DIA samples. MS^2^ spectra might also differ as fractionated DDA library spectra will not contain any co-fragmentation interferences that would occur in the original single-shot DIA data. ^24^ Nonetheless, we generated a hybrid library, by performing DirectDIA with the addition of DDA data which achieved greater proteome coverage as the library consisted of 124,908 precursors, 99,121 peptides and 8,285 protein groups **(Figure 2, 6).** All samples were run on identical analytical gradients and resulting thermal proteome profiling analysis performed in R **(Figure 2, 7)**.

### Protein Identification

There is a fundamental trade-off between achieving deep proteomic coverage and obtaining suitable throughput in proteomics studies, where chemoproteomics is no exception. The overall time required for each workflow was compared (**Figure 3A** and **Table S2A)**. In total, the thermal shift assay produced 48 samples (two treatment conditions: losmapimod and vehicle, each with eight temperatures and three biological replicates) that were either analysed by label-free single-shot DIA-MS or further labelled with TMT.

**Figure 3.**
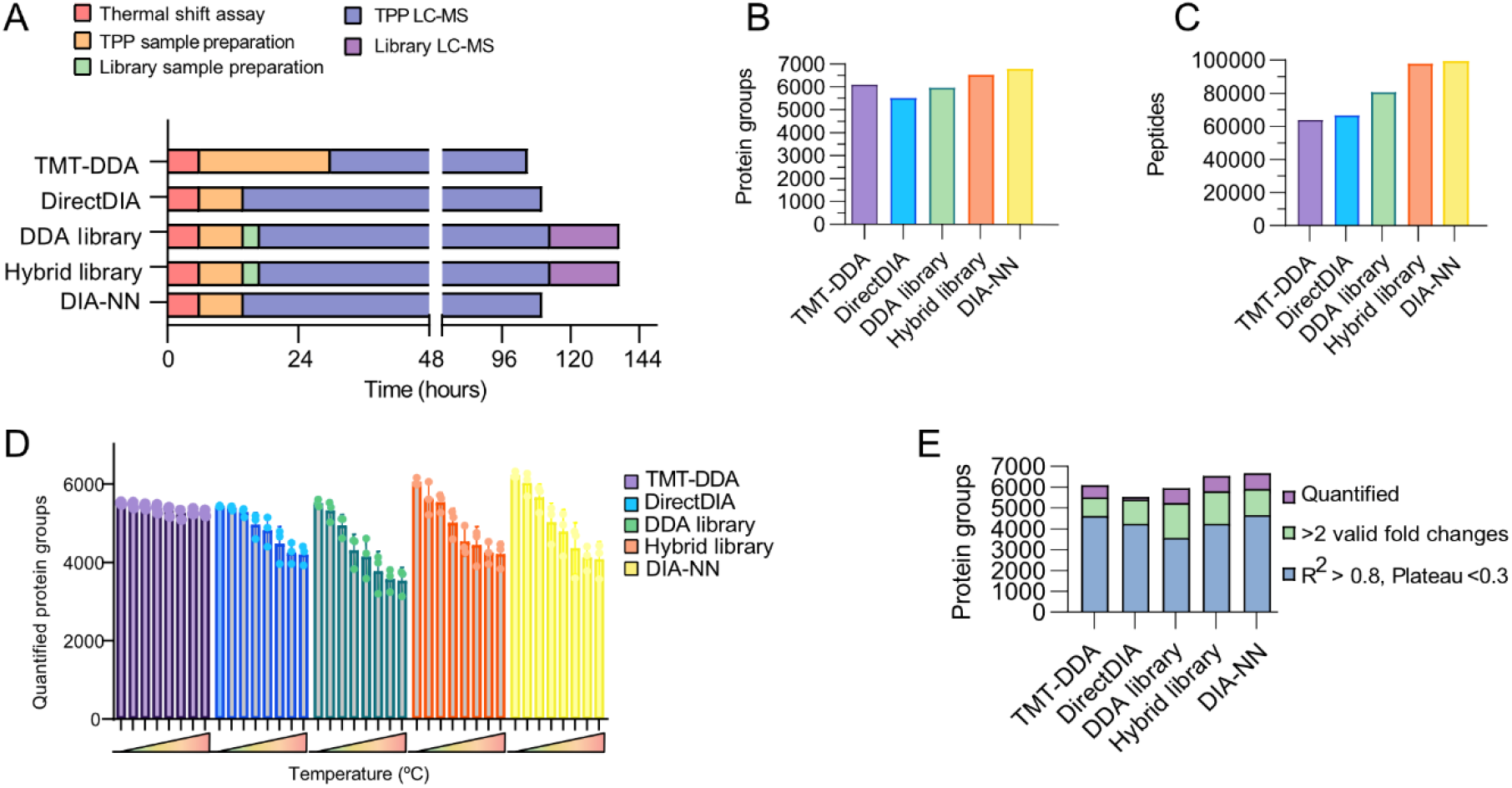
Performance overview of quantitative approaches for thermal proteome profiling. **A)** Bar plot showing the time required in hours for each method, comparing thermal shift assay (red), TPP-sample preparation (orange), library-sample preparation (green), instrument time for TPP-sample (blue) and library-sample (purple) analyses. **B)** proteins cumulative across all samples, **C)** number of peptides identified, **D)** the average number of quantified proteins across three biological replicates in each temperature soluble fraction and **E)** the average number of quantified proteins (purple), with two valid values for curve fitting (green) and of which sigmoidal curves were fit well (R^2^ > 0.8 and plateau < 0.3) (blue).

As shown in **Figure 3A**, the DIA workflows (including TPP sample preparation, any library-sample preparation, TPP-sample MS time or library acquisition MS time) consumed 110 h (Direct DIA and DIA-NN) and 137 h (DDA library and hybrid library). Cumulatively, TMT quantification required 105 h; 24 h for sample preparation, to label peptides of each biological replicate with a batch of 16-plex TMT, introducing additional clean-up steps whereas label-free single-shot DIA required only 8 h of sample preparation. Labelling efficiency checks added three hours of instrument time to the TMT experiment and would need to be fully processed and quality checked before sample preparation could proceed. The three sets of labelled samples were pooled, cleaned-up and fractionated taking 5 h, due to pools having larger sample volumes. This totalled 39 samples which altogether took 105 h of method time. Despite more sample handling time, TMT-DDA was the quickest method overall, with 75 h LC-MS time compared with 96 h LC-MS/MS analysis for the DIA TPP samples. Various factors may influence the decision of which DIA workflow to adopt, such as number of compounds/conditions and availability to instrument time, and so would need to be assessed on a project-specific basis. Nevertheless, the capacity for investigating the target landscape of multiple compounds, effects of combination therapies or resistance mechanisms becomes more feasible with a label-free approach or if within the same biological model, a project-specific library could be repurposed, becoming increasingly cost-effective.

Each approach identified a different number of cumulative protein identifications (**Figure 3B**; **Table S2B**). Overall, library-free DIA-NN yielded the greatest number of protein groups (6,669 protein groups) and peptide identifications (99,678 peptides), while DirectDIA data provided the lowest (5,518 protein groups; 66,612 peptides). TMT-DDA identified more protein groups (6,086 protein groups) compared to the DDA library analysis of the DIA data, likely reflecting the off-line fractionation, highlighting an advantage of the workflow and corroborating a previous benchmark comparison. ^11^ Although, the hybrid library (6,528 protein groups; 97,815 peptides) outperformed the DDA-library (5,954 protein groups; 80,630 peptides) as library performance is influenced directly by its quality and coverage, and the hybrid library consisting of both DDA, and DIA raw files was the most in-depth (**Figure 3C**). ^48^ Considerable overlap in identified protein groups was seen between all approaches, although less-so between DIA approaches and TMT-DDA (**Figure S2**). This finding highlights that DIA-based methods analyse slightly different portions of the proteome to DDA as on average 67 % of protein groups were common between DIA approaches and TMT-DDA, as previously reported.^49^ In DDA mode, the most abundant precursor peptide ions are isolated for acquisition of MS^2^ spectra, whereas the entire *m/z* range is selected in DIA, for an unbiased set of precursors, leading to variations in peptide, and ultimately protein identification. Moreover, peptides acquire a shift in mass and charge because of isobaric labelling, resulting in different peptides being fragmented.

### Melting Curve Fitting

Measuring significant shifts in protein thermal stability, indicative of compound-target engagement, relies on the successful construction and reproducible analysis of full protein melting curves. Therefore, the computational analysis of TPP datasets requires unique consideration. Raw abundance values from DDA or DIA analysis are quality filtered prior to curve fitting to exclude low-confidence identifications by number of peptide-spectrum matches (PSMs) or unique peptide number. As temperature increases, identification profiles become sparser by way of protein aggregation and the complexity of the soluble fractions becomes reduced. ^50^ TMT-DDA identified more protein groups in the less complex, higher-temperature samples than all DIA approaches (**Figure 3D**). However, data is required to have two valid fold-changes (and must therefore be detected in the lower reference temperature) for plotting the melting curve. Despite fewer missing values across the high-temperature profiles, several protein groups were not detected in the lower reference temperature for TMT-DDA resulting in 5,497 protein groups suitable for curve fitting. DIA-NN provided considerably more curves suitable for fitting (5,961 protein groups) along with the hybrid library analysis of DIA data (5,230 protein groups) (**Figure 3E, green**; and **Table S2B**).

Resulting sigmoidal curves were filtered for reliable protein melting point calculation prior to interpretation of the statistical and significance comparison **(Table S3-7)**. Only curves with a minimum coefficient of determination of < 0.8, indicating how well the fold changes fit the melting curve, and a plateau (lower horizontal asymptote) of < 0.3 were included, as recommended by Franken *et al.*^2^ TMT-DDA accurately quantified melting temperatures of 4,497 protein groups on average, performing comparably to DIA-NN generating an average of 4,458 melting curves (1 % less) (**Figure 2E, blue**). Interestingly, library-free mode with DIA-NN performed better than DirectDIA and hybrid library (both 6 % less relative to TMT-DDA). Finally, despite providing more protein identification than DirectDIA, DDA-library based analysis of the DIA dataset performed worst with a 20 % reduction in well-fit melting curves compared to DIA-NN and TMT-DDA, highlighting the significance of reliable quantification over number of protein identifications for TPP pipeline output. In fact, both library-free DIA approaches performed either equally or better than library-based DIA methods.

### Performance for target deconvolution

Results were filtered to include proteins with a significantly positive or negative fold change in melting temperature, of greater than 2°C, and a change in melting temperature (T_m_) standard deviation of less than two. As expected, the known target of losmapimod, MAPK14, was stabilised following treatment and detected by all five workflows (**Figure 4A-F**). On average, DIA approaches measured MAPK14 T_m_ to increase from 44.6 ± 0.1°C in vehicle controls, consistent with previous reports^15^, to 50.0 ± 0.3°C with losmapimod. This was independently validated by western blot (**Figure 4G**). Notably, the average T_m_ for MAPK14 after DMSO treatment measured by TMT-DDA was up to 0.9°C higher compared with DIA, possibly due to co-isolation of precursors reducing quantification accuracy, despite using the SPS-MS^3^ for acquisition; ^11^ i.e., an under measure of thermal shift could be an artefact to ratio compression. ^6^ DIA approaches measured a greater thermal shift suggests label-free DIA may be more sensitive to subtle changes in thermal stability.

**Figure 4.**
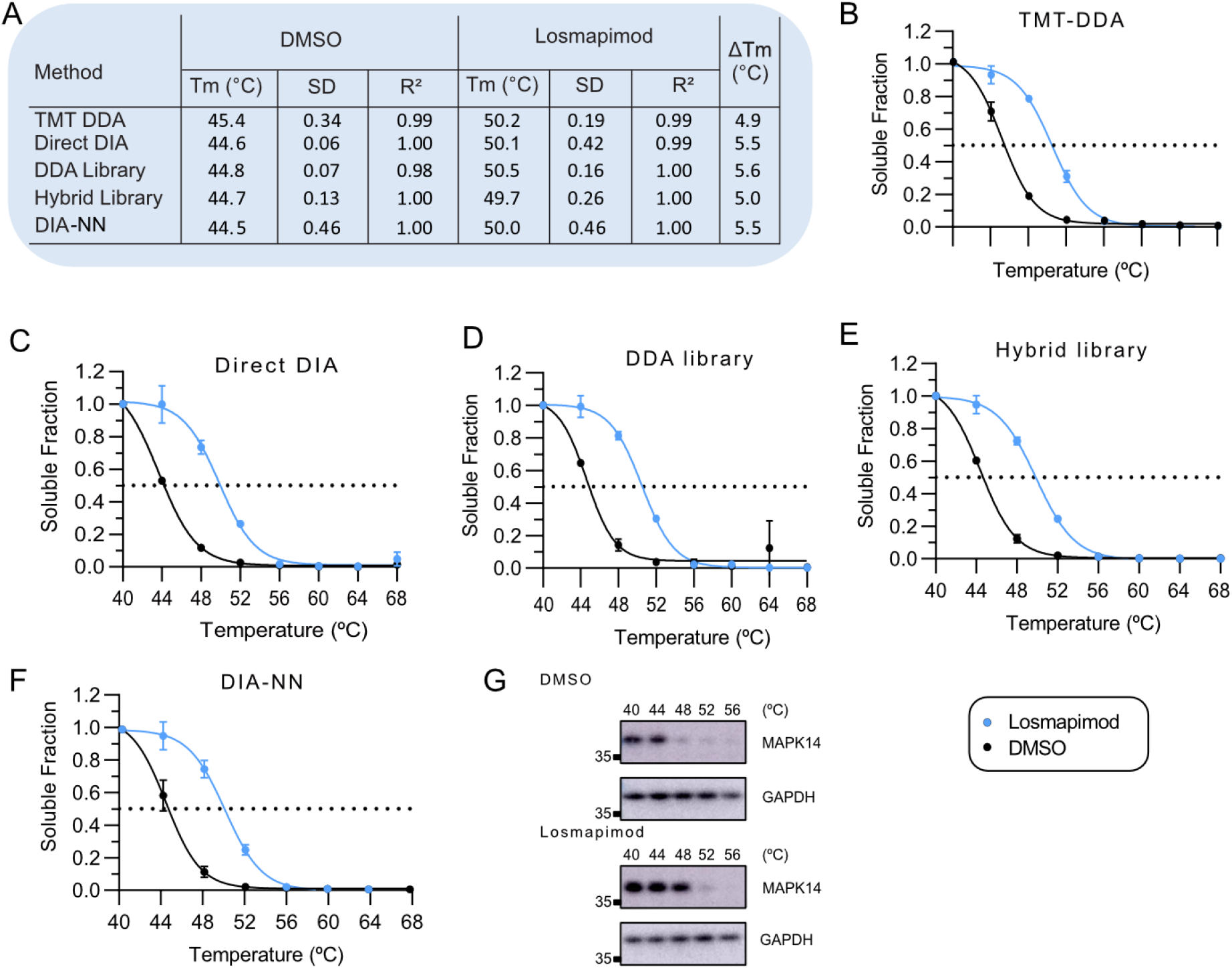
Melting curves for MAPK14. **A)** Table with significant shifts in melting temperature (*T_m_*) for MAPK14 quantified by **B)** TMT-DDA, **C)** Direct DIA, **D)** DDA library, **E)** hybrid library and **F)** library-free mode DIA-NN after treatment with losmapimod compared to vehicle. Error bars represent the SD of biological replicates. Dashed line indicates melting temperature (*T_m_*), where 50% of the protein is precipitated. **G)** Western blot analysis of the MAPK14 at the indicated temperatures in vehicle control (DMSO) and losmapimod treatment. GAPDH served as a loading control. Relative mobilities of reference proteins (masses in kDa) are shown on the left of each blot. SD, standard deviation; T_m_, melting temperature; R^2^ the coefficient of determination.

Given the therapeutic potential of losmapimod for the AML, we also explored its off-target landscape whilst evaluating the performance of all five pipelines. In addition to MAPK14, the downstream phosphorylation and interaction target MAPKAPK3 was significantly stabilised and detected by all methods. As seen in **Figure 5**, DIA-NN again detected a larger shift in melting temperature (+3.9 ± 0.1°C) in MAPKAPK3 following losmapimod treatment compared with TMT-DDA (+3.2 ± 0.1°C). Thermal stabilisation in this intact cell-based experiment may be due to inhibitor-induced biological changes in intracellular signalling.^6^ Phosphorylated proteins can display a different melting profile compared to their non-phosphorylated equivalents. ^6, 51^ To distinguish whether effects were primary (a consequence of direct drug binding) or secondary (a downstream cellular response to treatment), TPP in cell extracts without functioning PTM cellular machinery, would need to be performed.^4^ MAPKAPK3 stabilisation was independently validated by western blot (**Figure 5B)**.

**Figure 5.**
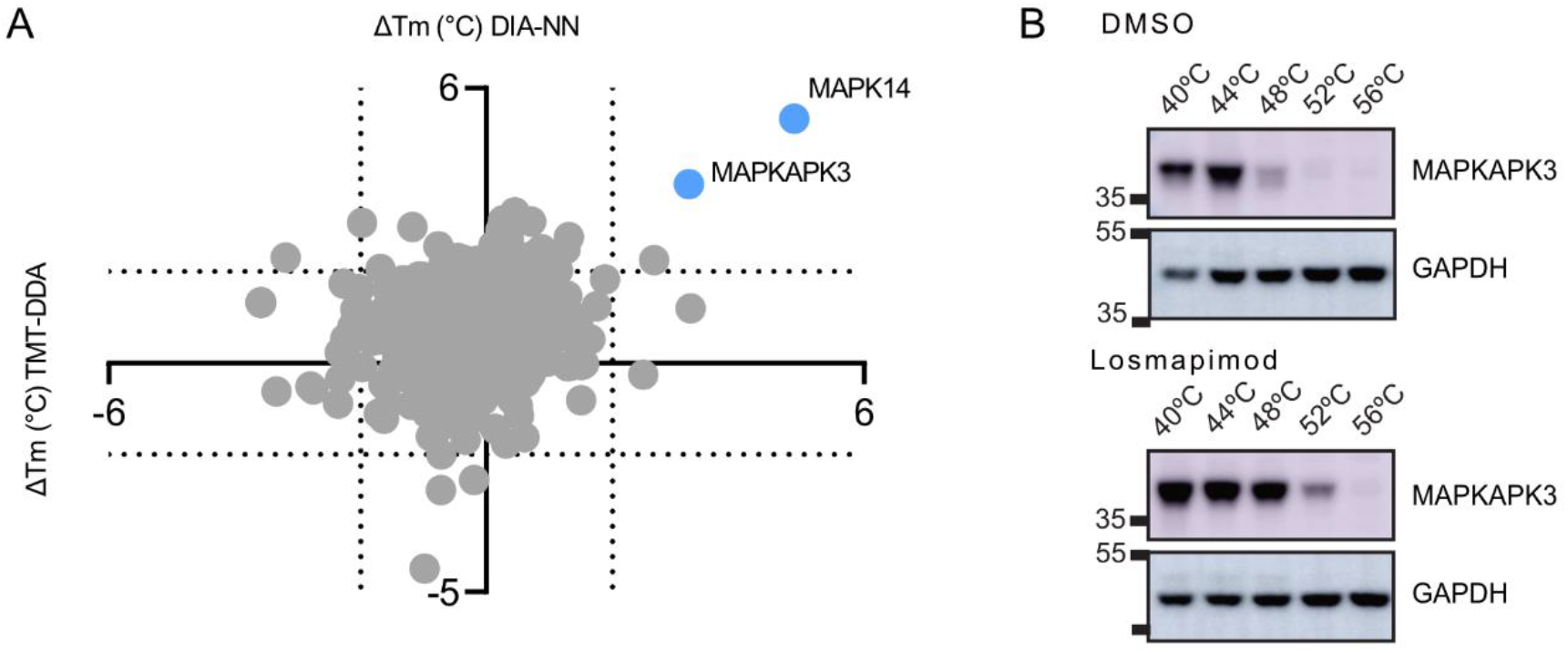
Target deconvolution of losmapimod by TMT-DDA and DIA analysis. **A)** Scatter plot of average T_m_ shifts calculated from well-fit sigmoidal curves of the three biological replicates following losmapimod treatment. Hits that were identified by both TMT-DDA and DIA-NN, which passed significance criteria are shown in blue. Proteins that did not pass significance criteria are shown in grey. **B)** Western blot analysis of the MAPKAPK3 at the indicated temperatures in vehicle control (DMSO) and losmapimod treatment. GAPDH served as a loading control. Relative mobilities of reference proteins (masses in kDa) are shown on the left of each blot.

Furthermore, TMT-DDA detected significant thermal stabilisation of myosin light chain kinase (MYLK) (3.6 ± 0.6°C) following treatment with losmapimod, but the protein was not detected in any DIA datasets (**Figure S3A**). A recent study reported MYLK to be a potentially new intracellular interaction partner of MAPK14^52^, which may have been stabilised by association due to MAPK14 target engagement. Stability changes in proteins present in a complex have been identified previously with TPP, for example, kinase complexes containing cyclins were stabilized by the kinase inhibitor staurosporine.^4^ TMT-DDA was also the only approach to detect significant thermal destabilisation of RAC-gamma serine/threonine-protein kinase (AKT3) with a −3.9 ± 1.7 °C shift in melting temperature from 53.6 ± 2.2°C to 49.7 ± 2.9°C following losmapimod treatment (**Figure S3B**). Of note, measured melting temperatures for AKT3 had great variability between biological replicates and if significance threshold is increased to > 0.01, the hit is removed. Although AKT3 was detected in the DIA data, its quantification did not pass the filtering criteria and was not used for statistical comparison, indicating TMT-DDA to be less stringent in identifying hits and more tolerant to noise. Overall, these results demonstrate the great specificity of losmapimod as a therapeutic agent, whilst demonstrating the ability of DIA to measure significant changes in thermal stability.

## Conclusion

In summary, we evaluated four DIA workflows that have not previously been compared for a TPP pipeline and benchmarked their performance with traditional TMT-DDA. This study highlights the potential of label-free DIA quantitative MS approaches for target deconvolution. Advantages were obtained with different DIA approaches, such as increased proteomic coverage by a library-based approaches, or improved throughput from library-free analysis, and in comparison to TMT-DDA, required less sample preparation time, but more instrument run-time, as expected with single-shot analyses. Ultimately, all methods compared in this study were unanimously able to identify MAPK14 as the primary target of the compound of interest, as well as detect MAPKAPK3 stabilisation, although smaller thermal shifts were observed by TMT-DDA compared to DIA, potentially a consequence of ion-interference or ratio compression. TMT-DDA identified two additional protein hits; one of which was thermally stabilised but not detected by DIA, which is a caveat of comparing different acquisition approaches. The second additional hit suffered from increased variability between replicates and was therefore filtered out in the DIA dataset suggesting TMT-DDA data may require a more stringent p-value threshold compared to DIA-NN. Comparison of Spectronaut and DIA-NN software for library-free DIA for TPP revealed differential performance as DIA-NN provided more well-fit melting curves, although software for the processing of DIA data is continually being developed and results may be different in alternative software versions. Nevertheless, given its superior performance and open-access nature, we advise DIA acquisition using library-free mode DIA-NN as a practical and cost-effective method for thermal proteome profiling.

## Author information

Corresponding authors

*(M.E.D) Maria Emilia Dueñas

*(J.L.M.-R.) José Luis Marín-Rubio

*(M.T.) Matthias Trost

## Authorship contributions

**Amy L. George**: Conceptualization, Formal analysis, Investigation of all TPP LC-MS/MS experiments, Methodology, Validation, Writing - Original Draft, Visualization. **Frances R. Sidgwick**: Methodology. **Jessica E. Watt:** Performed surface plasmon resonance. **Mathew P. Martin:** Performed surface plasmon resonance. **Matthias Trost:** Conceptualization, Resources, Writing - Review & Editing, Supervision, Funding acquisition. **José Luis Marín-Rubio:** Investigation of cell viability and proliferation, Validation, Writing - Review & Editing, Supervision, Project administration, Funding acquisition**. Maria Emilia Dueñas:** Methodology, Writing - Review & Editing, Supervision, Funding acquisition. All authors have read and agreed to the published version of the manuscript.

## Notes

The authors declare no competing financial interest.

## Acknowledgments

We would like to thank Abeer Dannoura for their technical support. This research was partly funded by a Wellcome Trust Investigator Award (215542/Z/19/Z). This research was partly funded by Newcastle Wellcome Trust Translational Partnership to J.L.M-R, M.E.D and M.T. M.E.D. is a Marie Sklodowska Curie Fellow within the European Union’s Horizon 2020 research and innovation programme under the Marie Skłodowska-Curie grant agreement No. 890296.

## Abbreviations

AGC: automated gain control
AKT3: RAC-gamma serine/threonine-protein kinase
AML: acute myeloid leukaemia
DDA: data dependent acquisition
DIA: data independent acquisition
FA: formic acid
FDR: false discovery rate
FDR: false discovery rate
FTMS: fourier transform mass spectrometry
GAPDH: glyceraldehyde 3-phosphate dehydrogenase
HCD: higher-energy collision-induced dissociation
IL: interleukin
LC-MS: liquid chromatography-mass spectrometry
LFQ: label-free quantification
m/z: mass-to-charge ratio
MAPK: mitogen-activated kinase
MAPKAPK: MAPK-activated protein kinase
MeCN: acetonitrile
MS: mass spectrometry
MS2: tandem mass spectrometry or MS/MS
MYLK: myosin light chain kinase
NP-40: Tergitol-type NP-40 and nonyl phenoxypolyethoxylethanol
PBS: phosphate buffered saline
RUs: response units
SD: standard deviation
SDS: sodium dodecyl sulfate
SDS-PAGE: sodium dodecyl sulfate–polyacrylamide gel electrophoresis
SPR: Surface plasmon resonance
SPS: synchronous precursor selection
TCEP: tris(2-carboxyethyl)phosphine
TEAB: triethylamonium bicarbonate
TFA: trifluoroacetic acid
TKIs: tyrosine kinase inhibitors
Tm: melting temperature
TMT: tandem mass tags
TNF-α: tumour necrosis factor alpha
TPP: thermal proteome profiling
UV: ultraviolet

## Supporting information

Figure S1. Losmapimod inhibits p38α and proliferation but not viability in AML cells; Figure S2. A Venn diagram comparing protein groups cumulatively identified between all approaches; Figure S3. Quantification of thermostability of A) MYLK3 and B) AKT, detected by TMT-DDA. Table S1. Summary of clinical trials of losmapimod. Table S2. A systematic comparison of the performance of five quantitative workflows for thermal proteome profiling. Table S3. TMT-DDA TPP analysis. Table S4. DirectDIA TPP analysis. Table S5. DDA library TPP analysis. Table S6. Hybrid library TPP analysis. Table S7. DIA-NN TPP analysis.

