## Supplemental Information for "A Comparison of Quantitative Mass Spectrometric Methods for Drug Target Identification by Thermal Proteome Profiling"

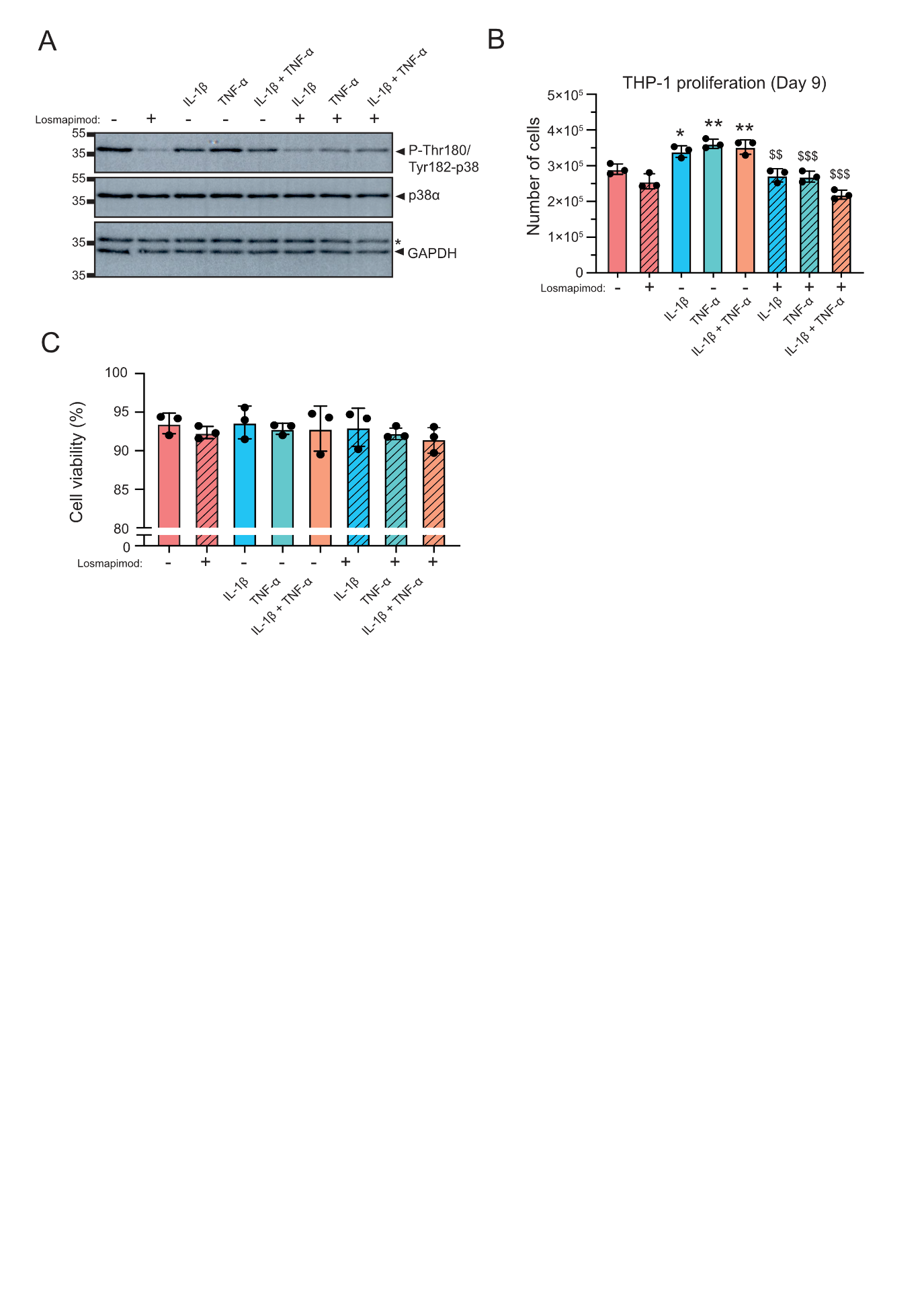

### Supplementary Figures

#### **Figure S1**. Losmapimod inhibits p38α and proliferation but not viability in AML cells.

A) Western blotting of the p38-MAPK phosphorylation in THP-1 cells pretreated with 1 µM losmapimod for 1 h before stimulation with 10 ng/mL IL-1β and/or 20 ng/mL TNF-α for 15 min. GAPDH serves as a loading control. A representative image with three biological replicates is shown. Relative mobilities of reference proteins (masses in kDa) are shown on the left the blot. B) Proliferation assay at day 9 in THP-1 cells with 1 µM losmapimod for 1 h before stimulation with 10 ng/mL IL-1β and/or 20 ng/mL TNF-α for 9 days. One-way ANOVA tests were performed (*, p<0.05; **, p<0.01; samples compared to untreated-sample without losmapimod; $$, p<0.01; $$$, p<0.001 samples compared to treated-samples without losmapimod). Error bars represent the standard deviation of three biological replicates. C) Cell viability assay at day 9 in THP-1 cells treated with 1 µM losmapimod for 1 h before stimulation with 10 ng/mL IL-1β and/or 20 ng/mL TNF-α. One-way ANOVA tests were performed. Error bars represent the standard deviation of three biological replicates.

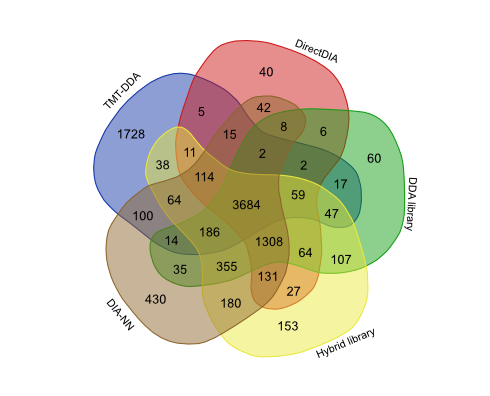

#### **Figure S2**. A Venn diagram comparing protein groups cumulatively identified between all approaches.

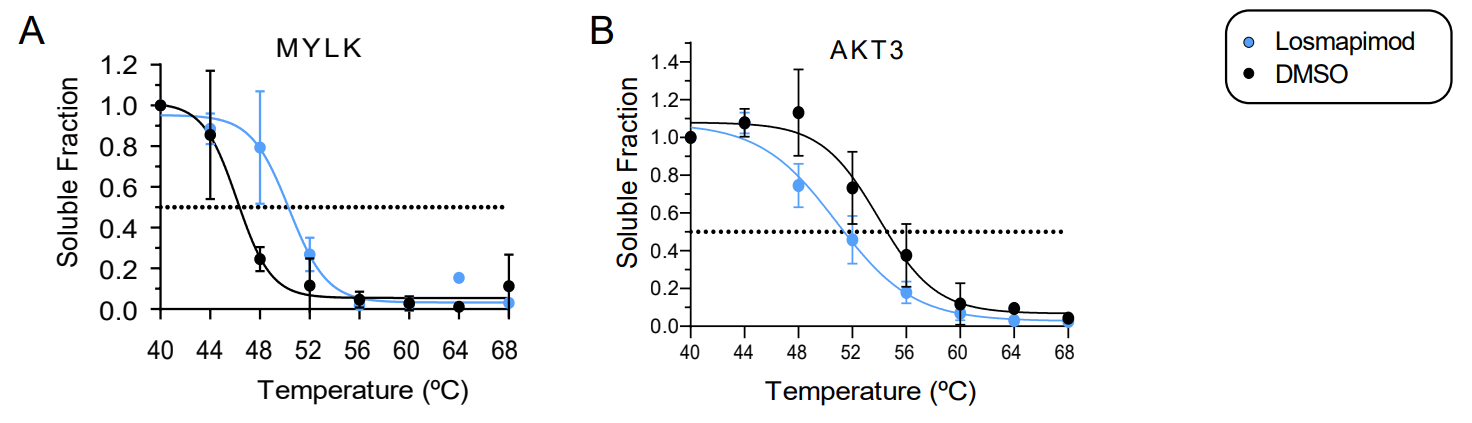

#### **Figure S3.** Quantification of thermostability of **A)** MYLK3 and **B)** AKT, detected by TMT-DDA.

Error bars represent the SD of three biological replicates.

### Supplementary Tables

#### **Table S1**. Summary of clinical trials of losmapimod.

A clinicaltrials.gov search on September 22, 2022, for losmapimod (keywords: losmapimod, GW856553, GW856553X, SB856553 or GSK-AHAB).

#### **Table S2**. A systematic comparison of the performance of five quantitative workflows for thermal proteome profiling.

**A)** The time required for each workflow, categorised by hours required for sample preparation, TPP analyses and any library generation. **B)** A breakdown of the total number of protein groups identified by each workflow, and results melting curves.

#### **Table S3.** TMT-DDA TPP analysis.

#### **Table S4.** DirectDIA TPP analysis.

#### **Table S5.** DDA library TPP analysis.

#### **Table S6.** Hybrid library TPP analysis.

#### **Table S7.** DIA-NN TPP analysis.
